# Distinct axo-protective and axo-destructive roles for Schwann cells after injury in a novel compartmentalised mouse myelinating coculture system

**DOI:** 10.1101/2023.05.19.541371

**Authors:** Clara Mutschler, Shaline V. Fazal, Nathalie Schumacher, Andrea Loreto, Michael P. Coleman, Peter Arthur-Farraj

## Abstract

Myelinating Schwann cell (SC)– dorsal root ganglion (DRG) neuron cocultures have been an important technique over the last four decades in understanding cell-cell signalling and interactions during peripheral nervous system (PNS) myelination, injury, and regeneration. While methods using rat SCs and rat DRG neurons are commonplace, there are no established protocols in the field describing the use of mouse SCs with mouse DRG neurons in dissociated myelinating cocultures. There is a great need for such a protocol as this would allow the use of cells from many different transgenic mouse lines. Here we describe a protocol to coculture dissociated mouse SCs and DRG neurons and induce robust myelination. Use of microfluidic chambers permits fluidic isolation for drug treatments, allows cultures to be axotomised to study injury responses, and cells can readily be transfected with lentiviruses to permit live imaging. We used this model to quantify the rate of degeneration after traumatic axotomy in the presence and absence of myelinating SCs and axon aligned SCs that were not induced to myelinate. We find that SCs, irrespective of myelination status, are axo-protective and delay axon degeneration early on. At later time points after injury, we use live imaging of cocultures to show that once axonal degeneration has commenced SCs break up, ingest, and clear axonal debris.

**Summary statement:** A novel compartmentalised dissociated mouse myelinating SC-DRG coculture system reveals distinct axo-protective and axo-destructive phases of Schwann cells on axon integrity after trauma.

## Introduction

Dissociated myelinating SC-DRG cocultures from rats were first developed by the Bunge laboratory in the 1980’s to investigate PNS myelination in a more dynamic way (Bunge *et al*, 1989; Eldridge *et al*, 1987). These cultures have been used to make seminal discoveries in uncovering the molecular mechanisms of SC myelination and responses of SCs and axons to injury (Taveggia *et al*, 2005; Guertin *et al*, 2005; Vaquié *et al*, 2019; Babetto *et al*, 2020; Salzer & Bunge, 1980). While there are available methodological descriptions for dissociated cocultures using rat SCs with either rat or mouse DRG neurons, there are no detailed published protocols using exclusively mouse cells (Taveggia & Bolino, 2018; Vaquié *et al*, 2018). The consensus within the field is that inducing myelination in dissociated mouse SCs is challenging. Certainly, induction of myelin differentiation with cyclic adenosine monophosphate (cAMP) analogues or elevating agents, such as forskolin, is more difficult in mouse SC monocultures compared to rat SC cultures. This is because mouse SCs require additional exogenous β-neuregulin-1 (βNRG1), plating on poly-L-lysine (PLL) instead of poly-D-lysine (PDL), and low concentration horse serum as opposed to foetal calf serum (Stevens *et al*, 1998; Arthur-Farraj *et al*, 2011). Indeed there has only ever been one laboratory detailing convincing myelin formation in dissociated mouse myelinating SC-DRG neuron cocultures (Stevens *et al*, 1998; Stevens & Fields, 2000). Protocols exist where endogenous or exogenous mouse SCs are used to myelinate DRG explant cultures (Päiväläinen *et al*, 2008; Stettner *et al*, 2013). Additionally, a more recent published protocol used endogenous SCs to repopulate dissociated mouse DRG axons, however SCs were only able to attain the pro-myelinating stage in coculture, further demonstrating the difficulty to get cultured mouse SCs to form mature myelin (Sundaram *et al*, 2021). Moreover, all the prior methods described using endogenous SCs or DRG explants preclude many experimental uses, such as using SCs from different transgenic animals, separate transfection of SCs and neurons with viruses for live imaging, and use of microfluidic chambers to allow injury studies and separate drug treatments to neurons or SCs. Furthermore, use of exogenous SCs in a DRG explant culture risks contamination from endogenous SCs and satellite glia migrating out of the DRG over time. This occurs because antimitotic treatment is unlikely to fully penetrate the whole DRG without prior dissociation and secondly, a compartmentalised culture system cannot be readily used with DRG explant cultures.

Recent studies using SC-DRG cocultures to investigate axon-SC interactions after injury have found conflicting results. A study using dissociated rat myelinating SC-DRG neuron cocultures in microfluidic chambers found that the presence of SCs accelerated the disintegration of axons after traumatic axotomy (Vaquié *et al*, 2019). In contrast, a study that seeded rat SC on mouse DRG axons, again in microfluidic chambers, but did not induce them to myelinate, found that the presence of SCs delayed axon degeneration (Babetto *et al*, 2020). Certainly data from *in vivo* studies, first in zebrafish and later in mouse have shown that SCs do participate in the breakup of the axon (Rosenberg *et al*, 2014; Villegas *et al*, 2012; Catenaccio *et al*, 2017; Vaquié *et al*, 2019). It is possible that the discrepancy between the two coculture studies is down to a species difference, given Babetto et al., 2020 combined rat and mouse cells whereas Vaquié et al., 2019 studied solely rat cells. Another explanation is that myelination status of the SCs may influence the outcome of the experiment as Vaquié et al., 2019 induced myelination prior to injury whereas Babetto et al., 2020 did not. Therefore, we decided to use our dissociated mouse SC - mouse DRG neuron coculture system to address the question of whether SCs accelerate or delay axon degeneration and whether the outcome depends upon myelination status.

Here we describe a detailed protocol for setting up dissociated mouse myelinating SC-DRG neuron cocultures in microfluidic chambers. We demonstrate how these compartmentalised cocultures can be used for performing axotomies, drug treatments, lentiviral infection of DRG neurons and SCs separately, and live imaging. SCs can be induced to robustly express myelin markers periaxin (PRX), myelin protein zero (MPZ) and myelin basic protein (MBP), and form electron dense myelin and nodal and paranodal structures. After axotomy, cocultured SCs replicate key parts of the *in vivo* injury response upregulating the major injury transcription factor, c-JUN (JUN), demyelinating and forming myelin ovoids (Arthur-Farraj *et al*, 2012; Arthur-Farraj & Coleman, 2021). We then show that after traumatic axotomy, SCs first have an axo-protective role, as severed DRG axons in the presence of SCs degenerate more slowly, compared to severed DRG axons cultured on their own. We find that both myelinating SCs and aligned SCs, which have not been induced to myelinate, are capable of delaying axon degeneration, suggesting that myelination status is not important in regulating this phenomenon. At later timepoints after axotomy, live imaging of cultures reveals that SCs help break up and ingest axonal fragments, clearing the debris.

## Results

### Establishing mouse dissociated myelinating SC-DRG neuron cocultures in microfluidic chambers

To establish dissociated mouse myelinating SC-DRG neuron cocultures, we dissected DRGs from E14 mice, enzyme dissociated these (see methods), and seeded the DRG neuronal cell suspension into the top compartment of microfluidic chambers on PLL and Matrigel^®^ coated Aclar^®^ coverslips (Fig. 1A). DRG neurons are then purified using the anti-mitotic cytosine Arabinoside (Ara-C) and are allowed to extend axons across a 150 μm microgroove barrier into an axonal compartment for up to seven days (Fig. 1B). Due to their size, DRG neurons cannot cross this microgroove barrier, and are exclusively present in the top compartment of the microfluidic chamber. By maintaining a hydrostatic pressure gradient between the two compartments, we were able to apply this anti-mitotic solely to the DRG compartment. P2-P4 neonatal sciatic nerves were then dissected, enzyme dissociated, and Ara-C purified in DMEM/5% horse serum (HS) for 72 hours to obtain cultured mouse SCs (Arthur-Farraj *et al*, 2011). Ara-C was withdrawn from microfluidic chambers for 48 hours before cultured mouse SCs were seeded into the axonal compartment of the chambers, where they were allowed to align with axons and proliferate for up to one week (Fig. 1C). SCs can then be induced to myelinate over the course of approximately three weeks through supplementation of their cell culture media with forskolin (10 μM), β neuregulin-1 (10 ng ml^−1^), Matrigel^®^ (1/100) and L-ascorbic acid (50 μg ml^−1^, Fig. 1C). Importantly, we found that L-ascorbic acid was insufficient to induce myelination alone, unlike in rat SC-DRG cocultures, and that the addition of Matrigel^®^ to the myelination medium was crucial in inducing robust myelination (data not shown). Forskolin and βNRG1 were included in the myelination medium as we have previously shown that these agents can induce myelin proteins in cultured mouse SCs (Arthur-Farraj *et al*, 2011). For a detailed step by step description of how to set up these cultures please see our methods section.

**Figure 1:**
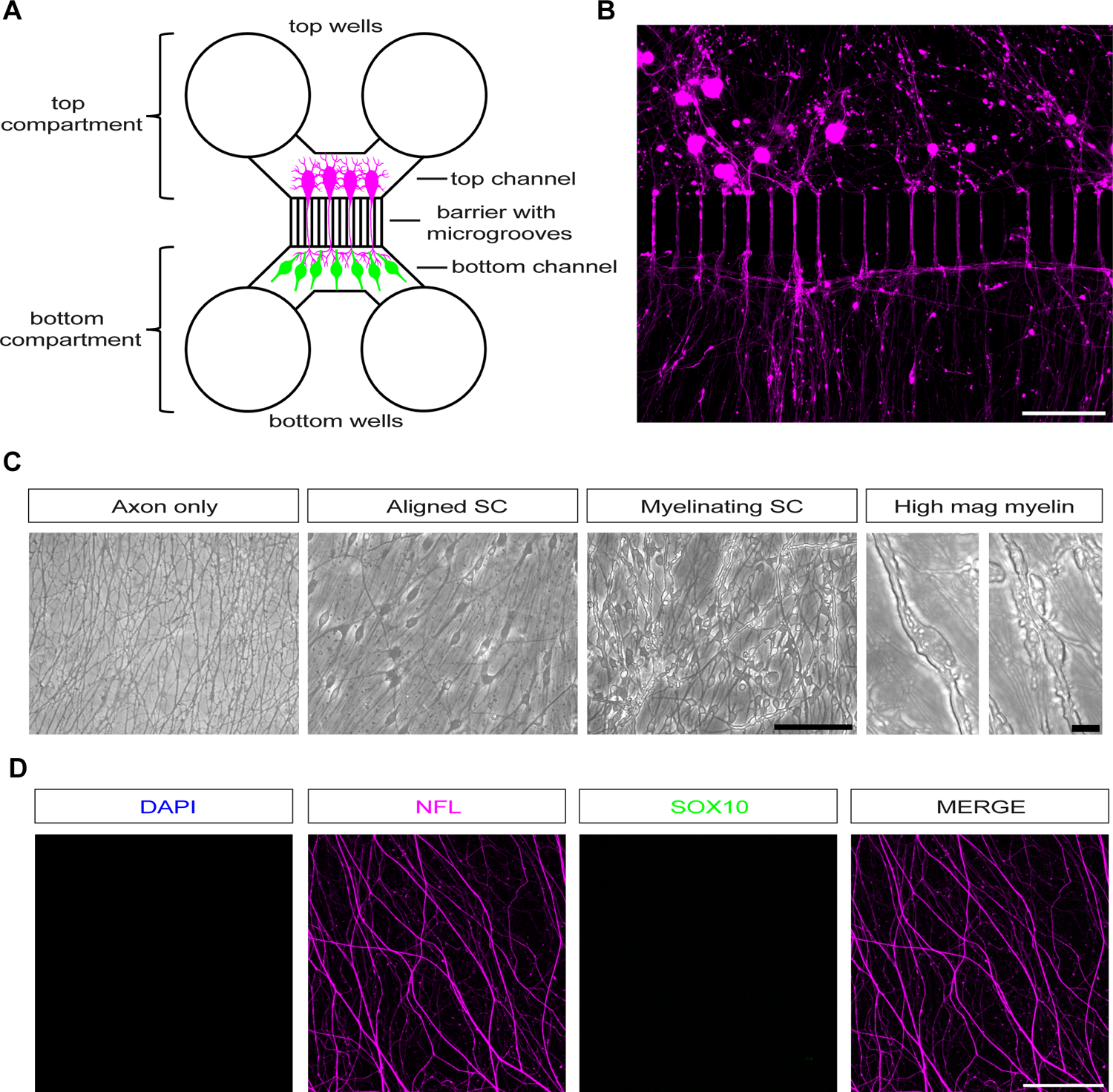
Murine Schwann cell dissociated dorsal root ganglion neuron cocultures in microfluidic chambers. **A.** Standard Neuron Device with a 150 μm microgroove barrier. Dissociated DRG neurons (magenta), which cannot cross the barrier due to their size, are seeded into the top channel and extend axons into the bottom channel, reaching the bottom wells. SCs (green) are then seeded in the bottom channel where they align with and myelinate axons. **B.** Dissociated DRG neurons (magenta) growing across the microgroove barrier. Scale bar 100 μm. **C.** Phase images of the bottom channel showing axons only, axons with aligned SCs (aligned SC), and axons with myelinating SCs (myelinating SC). Scale bar 100 μm. Higher magnification images showing myelin segments in cultures with myelinating SCs. Scale bar 10 μm. **D.** Confocal images of axon only cultures, showing axons (NFL, magenta), but no DAPI (blue) or SOX10 (green) signal in the axonal compartment of the microfluidic chamber. Scale bar 100 μm.

After approximately six weeks in culture, we were able to generate cultures with only axons in the axonal compartment, cultures with SCs aligned to axons but not induced to myelinate (aligned SC), and cultures with SCs myelinating axons (myelinating SC; Fig. 1C). Myelinating cocultures and cultures with aligned SCs can be easily distinguished by phase contrast microscopy, as myelin segments could be identified as long phase bright structures surrounding axons (Fig. 1C). Seven days of pulsed Ara-C treatment to the top compartment removed any non-neuronal cells. To confirm that axon only cultures are not contaminated by any potential surviving endogenous SCs that have migrated from the top compartment, we show no DAPI or SC specific SOX10 nuclear staining in the axonal compartment (Fig. 1D). In cocultures with aligned and myelinating SCs, we labelled SCs with the myelin associated protein PRX, which is expressed in the mouse sciatic nerve around birth, at the initiation of myelination (Gillespie *et al*, 1994). In cultures with aligned SCs, these cells have a much more diffuse, spread-out morphology, and a subset of these cells express PRX, while in myelinating cultures, strongly PRX positive myelin segments are present (Fig. 2A-C). To confirm that cocultured myelinated SCs formed compact myelin we performed electron microscopy (EM), which revealed compact myelin formation with multiple myelin wraps and formation of readily visible major dense (MDL) and intraperiod lines (IPL; Fig. 2D). Additionally, we immunolabelled cultures with antibodies to compact myelin proteins and found myelin sheaths were positive for MPZ and MBP (Fig. 2E and F). As myelinated fibres are organised into distinct domains, including the node of Ranvier, paranodal and juxtaparanodal regions, and the internode, we wanted to confirm if contactin associated protein 1 (CASPR1, also known as neurexin IV or paranodin) is confined to paranodal regions, where it normally accumulates in mature sheaths (Salzer, 2015; Einheber *et al*, 1997). We detected CASPR1 protein in the correct distribution, labelling the paranodal region, adjacent to regions of MPZ labelling (indicating compact myelin) in our myelinating SC-DRG neurons cocultures (Fig. 2G).

**Figure 2:**
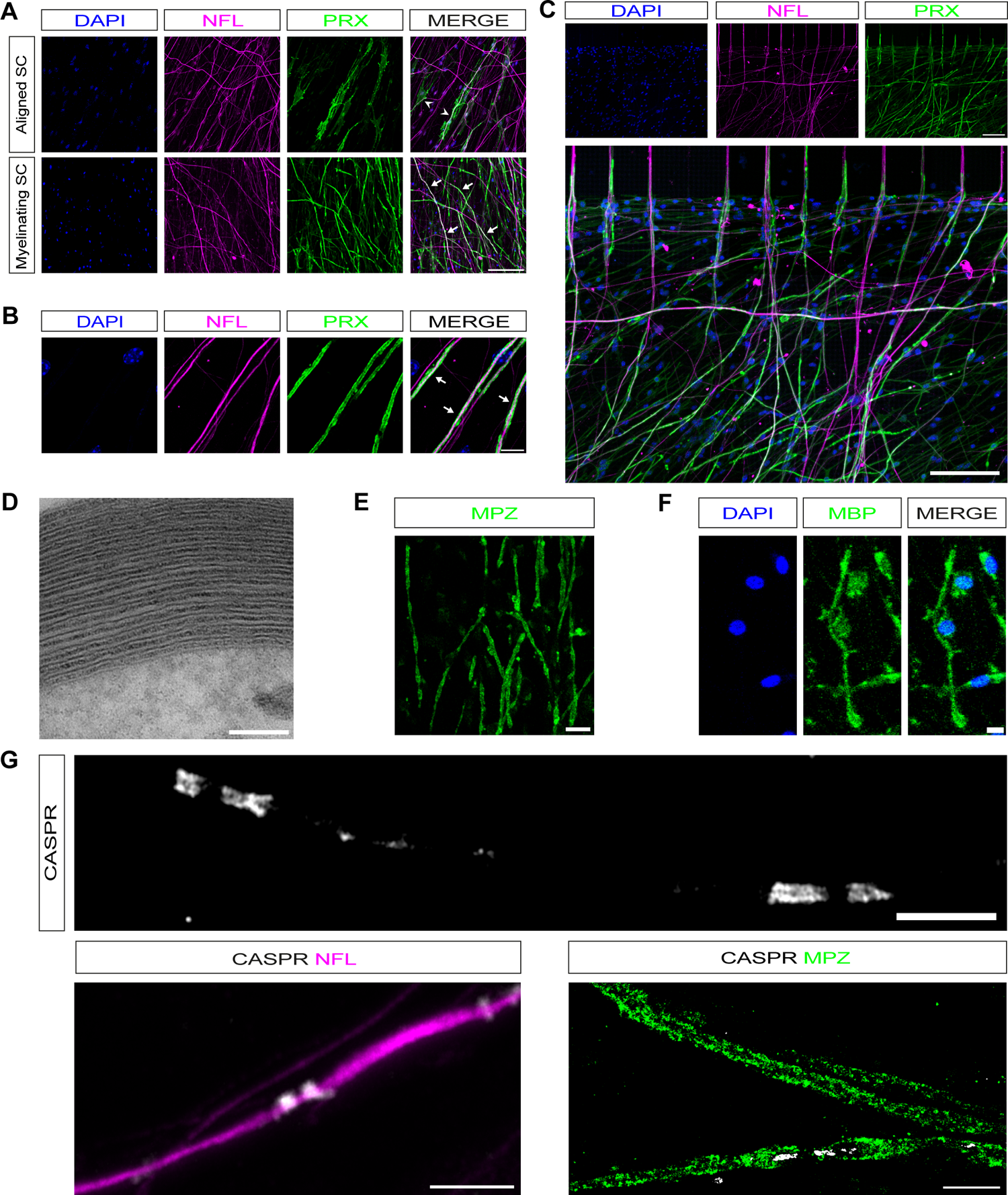
Schwann cells in myelinating cocultures can be induced to myelinate. **A.** Confocal images of dissociated DRG neuron cocultures with aligned SC or myelinating SC. Axons are NFL-labelled (magenta), and SCs PRX-labelled (green). Aligned SCs have a more diffuse morphology (white arrowheads), while myelin segments are present in myelinating SCs (white arrows). Scale bar 100 μm. **B.** Higher magnification images showing NFL-labelled axons (magenta) covered by myelin segments (PRX, green, white arrows). Scale bar 10 μm. **C.** DRG neurons (NFL, magenta) growing across the barrier in chambers with SCs (PRX, green) that have been induced to myelinate. Scale bar 100 μm. **D.** Electron micrograph of electron dense myelin in cocultures with myelinating SCs. Scale bar 100 nm. **E.** MPZ (green) labelled myelinating SCs. Scale bar 20 μm. **F.** MBP (green) labelled myelinating SCs. Scale bar 10 μm. **G.** In mature myelinating cultures, SCs CASPR (white) can be detected in the characteristic staining pattern marking paranodes. Scale bar 5 μm. Paranodal CASPR co-localised on axons (magenta). Scale bar 5 μm. When colabelling with MPZ (green), CASPR is localised to paranodal loops adjacent to a Node of Ranvier. Scale bar 10 μm.

In summary our dissociated mouse myelinating SC-DRG neuron cocultures develop robust compact myelin, as evidenced by myelin protein immunostaining and EM. Furthermore, our myelinating cocultures develop nodal/paranodal structures similar to cocultures prepared with rat SCs and DRG neurons.

### Axotomy in SC-DRG neuron cocultures replicates characteristic axonal and SC injury responses

After injury, SCs transform into repair SCs, which are characterised by a strong upregulation of the transcription factor JUN, and myelin breakdown by SCs through a process termed myelinophagy (Jessen & Arthur-Farraj, 2019; Gomez-Sanchez *et al*, 2015; Arthur-Farraj *et al*, 2012; Parkinson *et al*, 2008). To test whether axons and SCs respond to injury in a similar way in our coculture model as they do *in vivo*, we performed axotomies on cultures, using a scalpel under a light microscope, after carefully removing the chamber. In myelinating SC-DRG cocultures, 12 hours after axotomy, we fixed and immunolabelled cultures for Neurofilament light chain (NFL) and found that many axons distal to the site of axotomy had started to degenerate (n=4; Fig. 3A). Additionally, we noted a strong upregulation of JUN protein in SCs 12 hours after axotomy (Fig. 3B). Fluoromyelin labelling 48 hours post axotomy demonstrated myelin-ovoid formation, suggestive of active SC demyelination (Jung *et al*, 2011); Fig. 3C). We confirmed SC demyelination in our axotomised cocultures using electron microscopy, identifying characteristic demyelinated profiles surrounding degenerated axons, similar to what has been previously shown in rat SC-DRG cocultures (Fernandez-Valle *et al*, 1995) Fig. 3D).

**Figure 3:**
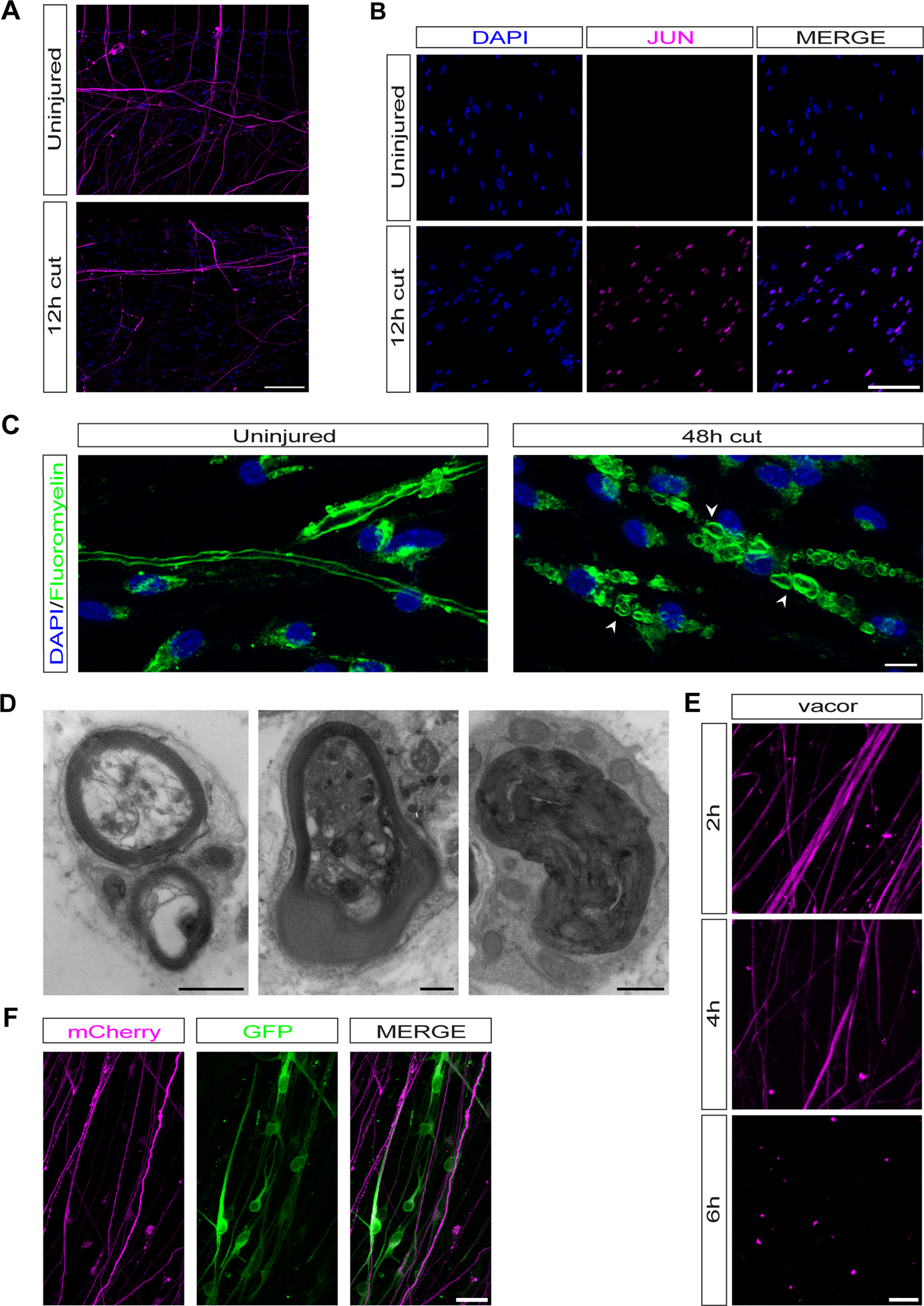
SC-DRG neuron cocultures replicate characteristic axonal and Schwann cell injury responses after axotomy. **A.** 12 hours post axotomy many of the axons (magenta) in SC-DRG neuron cocultures have degenerated. Scale bar 100 μm. **B.** SC JUN in uninjured cultures and 12 hours post axotomy. With identical imaging conditions, a signal can only be detected 12 hours post axotomy. Scale bar 100 μm. **C.** Myelinating SCs demyelinate after extended periods of time (48 hours) after axotomy. Myelin ovoids and myelin debris (both identified by white arrowheads) are present in fluoromyelin (green) labelled cultures. Scale bar 10 μm. **D.** Electron micrographs of myelinating SCs 48 hours post axotomy showing characteristic demyelinated profiles surrounding degenerated axons. Scale bar 1 μm. **E.** Vacor induces neurodegeneration when applied to the top compartment of cocultures. NFL (magenta). Scale bar 10 μm. **F,** Neurons and SCs can be lentivirally infected prior to plating in microfluidic chambers. Neurons infected with LV-CMV-mCherry (MOI 2, magenta) and SCs with LV-CMV-GFP (MOI 200, green). Scale bar 20 μm.

In addition to traumatic axotomy, our cocultures can also be used for drug treatments. To demonstrate this, we treated only DRG cell bodies with the specific SARM1 agonist vacor, which induces specific degeneration of axons and neuronal cell bodies but not SCs, which are completely insensitive to SARM1 agonists (Loreto *et al*, 2021; Fazal *et al*, 2023). 50 μM vacor addition to DRG cell bodies induced axon degeneration within 6-8 hours in our cocultures (n=4, Fig. 3E). Finally, we also demonstrate that DRG neurons and SCs can be separately infected with lentiviruses (LV) to permit live cell imaging. Here we infected mouse DRGs directly after the DRG dissociation step with an mCherry expressing LV (LV-CMV-mCherry) and mouse SCs were infected with a GFP expressing LV (LV-CMV-GFP) after Ara-C purification and prior to seeding in the microfluidic chamber (Fig. 3F). We found both mouse DRGs and SCs transfected best with LVs in suspension with slow centrifugation (see methods). Interestingly, we found that embryonic DRG neurons were almost completely resistant to LV transfection if they had already been cultured for 2-3 days (data not shown). In summary, our mouse myelinating SC-DRG cocultures can be axotomised to study SC-axon interactions as they reliably replicate *in vivo* cell behaviours. Furthermore, these cocultures can be used to study drug induced neurodegeneration and both DRG neurons and SCs can be separately transfected with LVs for live imaging and genetic disruption studies.

### Schwann cells are axo-protective at early timepoints and axo-destructive at late timepoints after axotomy

Recent evidence has shown both axo-protective and axo-destructive roles for SCs in cocultures. Two possible explanations for the discrepancy in results is that Babetto et al., 2020 used axotomy in rat SCs seeded on mouse DRG axons without inducing myelination, whereas Vaquié et al., 2019 studied axotomy in myelinating rat SC-DRG cocultures (Vaquié *et al*, 2019; Babetto *et al*, 2020). To examine whether myelination status influences the axon degeneration rate *in vitro*, we performed axotomies on axon only cultures, cultures with aligned SCs and cultures with myelinating SCs. Cocultures were fixed at three, six, nine, and 12-hours post axotomy and immunolabelled with NFL to assay axonal integrity (Fig 4A). Importantly, we found that fixed myelinated cultures needed to be permeabilised with acetone to allow full penetration of NFL antibodies to label axons in their entirety through myelinated segments (data not shown). While there was negligible degeneration in all cultures at three hours post axotomy, there was noticeably more degeneration at six hours post axotomy in axon only cultures (29.00 ± 1.64%, n=5), compared to both aligned SC, and myelinating SC cocultures (8.19 ± 2.57%, n=3 p<0.0001, and 8.14 ± 0.65%, n=4 respectively, p<0.0001). Both aligned and myelinating SC cocultures also showed significantly lower amounts of degeneration at both nine and 12 hours post axotomy in comparison to axon only cultures (axon only: 9h, 47.39 ± 1.34%, n=3, and 12h, 76.68 ± 6.60%, n=3; aligned SC: 9h, 30.05 ± 3.05%, n=4, p=0.0004 and 12h, 44.61 ± 4.72%, n=3, p<0.0001; myelinating SC: 9h, 33.12 ± 0.61%, n=3, p=0.0059 and 12h, 48.38 ± 7.75%, n=3, p<0.0001) (Fig. 4B). Between aligned and myelinating SC cocultures, there were no significant differences in amounts of axon degeneration (3h: p=0.65, 6h: p=0.98, 9h: p=0.44, 12h: p=0.70).

**Figure 4:**
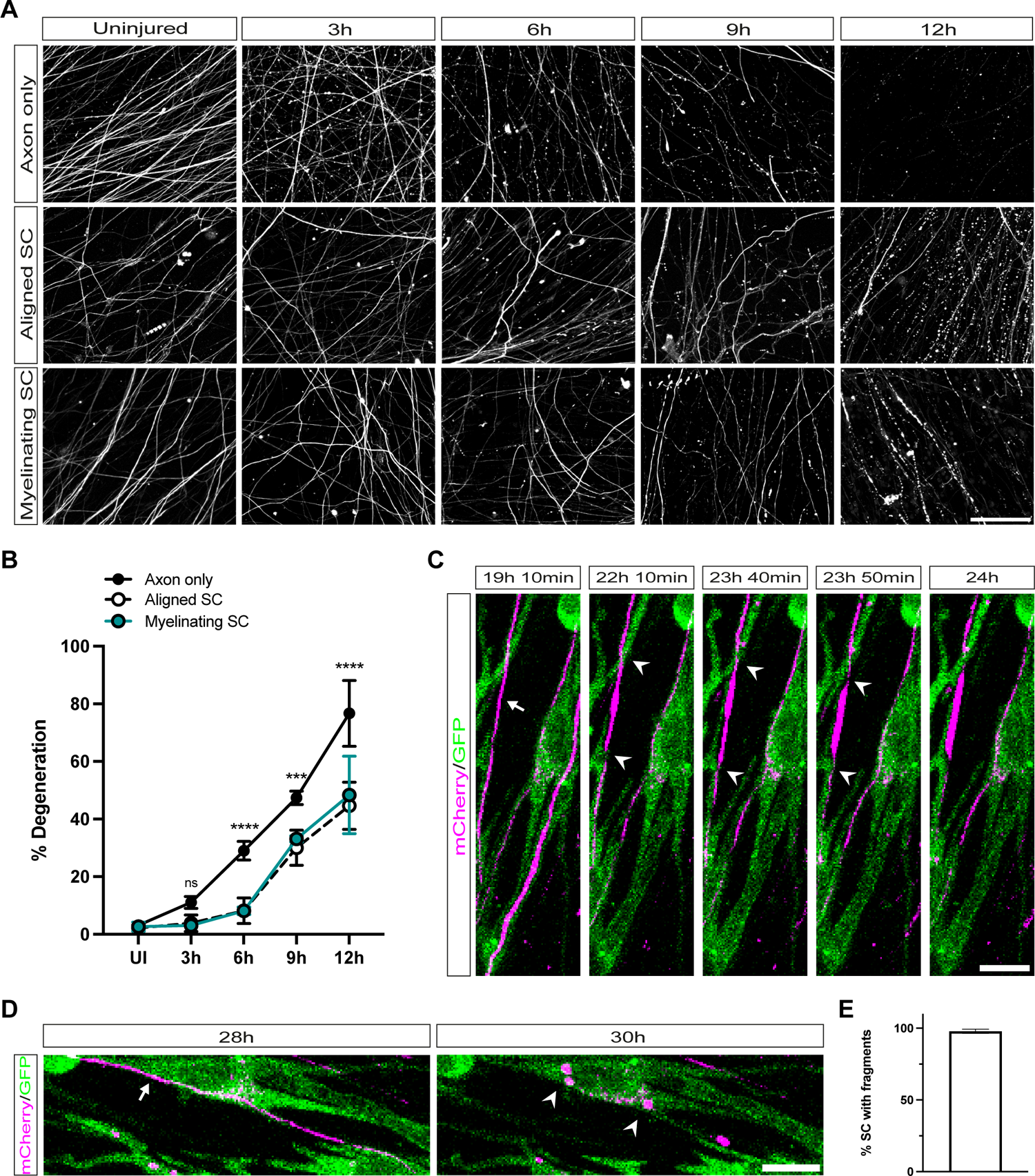
Schwann cells are both axo-protective and axo-destructive after axotomy. **A.** Confocal images of NFL-labelled cocultures prior to axotomy at the microgroove barrier (uninjured), and 3, 6, 9, and 12 hours after axotomy. Axon only cultures show earlier signs of degeneration than cultures with SCs. Scale bar 100 μm. **B.** Quantification of axon degeneration after axotomy. Only axon only cultures show statistically substantial degeneration at 6 hours post axotomy (29 ± 1.44%, n=5). In axon only cultures this degeneration increases to 47.39 ± 1.34% at 9 hours post axotomy (n=3), and finally 76.68 ± 6.60% at 12 hours post axotomy (n=3). Cocultures with SCs show little degeneration at 3 hours post axotomy (aligned SC: 4.87 ± 1.65%, n=3; myelinating SC: 3.048 ± 0.06%, n=3) and 6 hours post axotomy (myelinating SC: 8.14 ± 0.65%, n=4; aligned SC: 9.19 ± 2.57%, n=3). At 9 and 12 hours post axotomy, axons associated with both aligned and myelinating SCs start to degenerate (aligned SC: 9h, 30.05 ± 3.05%, n=4 and 12h, 44.61 ± 4.72%, n=3; myelinating SC: 9h, 33.12 ± 0.61%, n=3 and 12h, 48.38 ± 7.75%, n=3). There were no significant differences in axon degeneration rates between aligned and myelinating SCs cocultures (3h: p=0.65, 6h: p=0.98, 9h: p=0.44, 12h: p=0.70). Axon only cultures show significant differences to cultures with aligned or myelinating SCs at 6 (aligned SC: p<0.0001, myelinating SC: p<0.0001), 9 (aligned SC: p=0.0004, myelinating SC: p=0.0059), and 12 (aligned SC: p<0.0001, myelinating SC: p<0.0001) hours post axotomy. Results shown as Mean ± SEM. Statistical significance determined by 2-way ANOVA. Statistical significance for comparisons between axon only cultures and aligned SC cocultures are displayed on the graph. **p<0.01. ***p<0.001. ****p<0.0001. **C.** Confocal images of mCherry-labelled axons (magenta) and GFP-labelled SCs (green). At an early timepoint (19 h 10 min) an intact axon is visible (white arrow). 3 hours later (22 h 10 min), the axon is starting to be constricted (white arrowheads). This continues until 23 h 50 min after axotomy, when constrictions are clearly visible, and the axon is swollen between two constrictions. 24 hours after axotomy, the axon then breaks apart. Scale bar 10 μm. **D.** Confocal images of mCherry-labelled axons (magenta) and GFP-labelled SCs (green) with intact axons (white arrow, 28 h post axotomy) just before axon degeneration and with mCherry fragments (white arrowheads) within SCs 30 h post axotomy, once axon degeneration has occurred. Scale bar 10 μm. **E.** Quantification of number of SCs with fragments in the first 48 h post axotomy. s97.84 ± 1.462 %, n=10. Data represented as mean ± SEM.

Since Vaquié et al., 2019 showed SCs appear to accelerate axon degeneration in rat cocultures at later timepoints after axotomy, we were interested to see whether SCs in our cocultures help break up, ingest, and clear axonal fragments. We used cocultures where axons were labelled with LV-CMV-mCherry and SCs with LV-CMV-GFP and we live imaged cultures for up to 48 hours post axotomy. In these cocultures, we visualised axons break into large fragments surrounded by GFP positive SC processes, similar to the constricting actin spheres described by Vaquié et al., 2019. We then visualised SCs phagocytose and digest mCherry-labelled axonal fragments (Fig. 4C; supplemental video 1). When we quantified this phenomenon, we found that 97.84 ± 1.462% of SCs in our cocultures contained mCherry-labelled axonal fragments (Fig. 4D).

Thus, in a dissociated mouse coculture system, the presence of SCs delays the onset of axon degeneration at sites distal from the axotomy. Furthermore, this delay in degeneration does not appear to be reliant on the myelination status of the SCs. At later timepoints, once axons start to degenerate, SCs fragment, ingest and clear axonal debris.

## DISCUSSION

Here we describe a protocol to set up dissociated mouse myelinating SC-DRG neuron cocultures in microfluidic chambers. These can be utilised to study myelination and injury responses and can be adapted for the application of drugs to different cellular compartments and for live imaging. Furthermore, as this is a purely mouse cell culture system, one can utilise the vast array of transgenic and knockout lines available to study neuron-SC interactions in more detail. Our cultures display several hallmarks of mature myelin sheaths, with electron dense myelin, compact myelin proteins and a CASPR immunolabelling patterning suggesting assembly of paranodal and nodal structures. Axotomy of our cocultures faithfully replicates several of the key cellular events seen after nerve injury, including axon degeneration, upregulation of the key injury transcription factor c-JUN in SCs and SC demyelination with myelin ovoid formation.

Achieving myelin differentiation of mouse SCs, whether in monocultures or in coculture, has historically been very difficult. There are very few published reports of robust myelination in dissociated mouse SC-DRG neuron cocultures or of myelin differentiation in mouse monocultures (Stevens *et al*, 1998; Arthur-Farraj *et al*, 2011). Furthermore, there are no detailed published protocols for either. The majority of the field use rat SCs for *in vitro* myelination studies as it is much easier to achieve myelin differentiation and there are a number of published protocols (Eldridge *et al*, 1987; Vaquié *et al*, 2018; Taveggia & Bolino, 2018; Monje, 2020; Morgan *et al*, 1991). However, using our protocol it is possible to set up dissociated mouse myelinating SC-DRG neuron cocultures and take advantage of the ability to use cells from various transgenic mouse lines to study axon-SC interactions during myelination, injury, and regeneration. Furthermore, our protocol is complementary to the recently described mouse myelinating SC-motor neuron coculture system (Hyung *et al*, 2021). To date, there have been no published studies of successful myelination in a human SC-neuron coculture systems. Despite this rat SCs have been shown to readily myelinate human-induced programmable stem cell (iPSC)-derived sensory neurons and an iPSC-derived peripheral nerve organoid system which does contain myelinating SCs has recently been described (Clark *et al*, 2017; Van Lent *et al*, 2022).

We used our cocultures to test if the presence of SCs was predominantly axo-destructive or axo-protective after injury. We have identified two distinct phases of SC-axonal interaction post axotomy in coculture. At timepoints up to 12 hours post axotomy we observe that SCs delay the initiation of axon degeneration, however once the axon degenerates, live imaging up to 48 hours post axotomy demonstrates that SCs help break axons into large fragments and then ingest and clear axonal debris. These findings confirm and move to reconcile the seemingly contrasting observations of both Babetto et al., 2020 and Vaquié et al., 2019. (Vaquié *et al*, 2019; Babetto *et al*, 2020). The discrepancy in findings between our study and the two previous studies is unlikely to be a species difference given that both prior studies used rat SCs, and we have used exclusively cells from mice. Additionally, we have shown that the axo-protective observation does not rely on myelination status, which was another difference between the two original studies. A third key difference was that ourselves and Babetto et al., 2020 quantified the degeneration rates of the very distal neurites, in keeping with assays on axon degeneration performed in solely neuronal cultures (Loreto & Gilley, 2020). The study by Vaiquié et al., 2019 quantified axon fragmentation proximally in the microgrooves at timepoints starting from 12 hours after axotomy. This is a timepoint after we had seen degeneration distally in all of our cultures, consistent with published timescales for axon degeneration in mouse DRG neuron monocultures (Sasaki *et al*, 2009). It is possible that imaging proximally at later timepoints reduced the potential of identifying a subtle axo-protective phenotype but clearly displayed a role for SCs in the fragmentation and clearance of axonal debris (Vaquié *et al*, 2019).

Multiple independent findings from *in vivo* studies have demonstrated that SCs help break up the axon during or slightly after programmed axonal degeneration is initiated during nerve trauma (Catenaccio *et al*, 2017; Rosenberg *et al*, 2014; Villegas *et al*, 2012; Vaquié *et al*, 2019). We have also confirmed this phenomenon in a 2-photon axotomy model in zebrafish larvae (P. Arthur-Farraj unpublished observation). More recently Babetto et al., 2020 showed that SCs upregulate glycolysis within the first two days after traumatic nerve injury and this has an axo-protective effect *in vivo* (Babetto *et al*, 2020). SCs have been shown to promote axonal and neuronal survival in other situations, including during axon regeneration in both the acute and chronic setting, old age and in neuropathy (Arthur-Farraj *et al*, 2012; Wagstaff *et al*, 2021; Hantke *et al*, 2014; Fontana *et al*, 2012; Painter *et al*, 2014). It is thus likely that SCs have both axo-protective and axo-destructive roles, *in vivo*, after traumatic nerve injury. Future studies will be needed to detail the precise *in vivo* timing of these different cellular phases of SCs on axon integrity after nerve trauma.

In summary, we have described a detailed method of setting up dissociated mouse myelinating SC-DRG neuron cocultures in microfluidic chambers. These cultures can be transfected, readily live imaged, used for studying myelination and cellular responses to injury and regeneration, and used for drug studies. Most importantly SCs and DRG neurons from various transgenic mice can be used to perform *in vitro* analysis to complement findings from *in vivo* transgenic mouse studies.

## Methods

### Animals

All research complied with the Animals (Scientific Procedures) Act 1986 and the University of Cambridge Animal Welfare and Ethical Review Body (AWERB). Wild-type C57BL/6J mice were obtained from Charles River Laboratories and were held under standard specific pathogen free conditions.

### Coculture Axotomy

Traumatic axotomies were carried out by carefully removing the microfluidic chamber from the Aclar^®^ coverslip using sterile forceps and severing axons with a surgical blade under a light microscope. Axotomies were carried out at the level of the microgroove barrier. To confirm all axons were severed, a second higher cut was performed and axons between the cut sites removed using the surgical blade. Vacor axotomies were carried out by media changing the top compartment to media supplemented with 50 μM vacor (Greyhound Chromatography - N-13738).

### Quantification of Degeneration

Five images were quantified per culture, taken in comparable locations in each culture. A line was drawn across each image, and each axon crossing this line was either scored as degenerated or intact. Images were blinded prior to quantification.

### Immunocytochemistry

Cells were fixed for 10 minutes in 4% paraformaldehyde (PFA; Electron Microscopy Sciences) diluted in phosphate buffered saline (PBS) at RT. Axon only cultures were permeabilised in PBS + 0.5% Triton (Merck) + 5% HS + 5% donkey serum (DS, Merck - D9663) at RT for 1 hour. Cocultures with SCs were permeabilised in 50% Acetone for 2 minutes, 100% Acetone for 2 minutes, 50% Acetone for 2 minutes (all at RT), and then blocked in PBS + 0.5% Triton + 5% HS + 5% DS at RT for 1 hour. Myelinating cocultures stained for MBP, MPZ or CASPR were permeabilised in 50% Acetone for 2 minutes, 100% Acetone for 2 minutes, 50% Acetone for 2 minutes (all at RT), 100% Methanol at −20°C for 10 minutes, and then blocked in PBS + 5% HS + 5% DS at RT for 1 hour. Cultures were immunolabelled by incubating overnight at 4°C with primary antibodies. Primary antibodies were visualised using Alexa 488-, 568, and 647-conjugated secondary antibodies. DAPI (4’,6-diamidino-2-phenylindole, Thermo Fisher Scientific - D1306) was used 1:10000. Cells were mounted using Citifluor Glycerol Pbs Solution AF1 (Agar Scientific Ltd) and sealed using nail varnish. For confocal imaging a Zeiss LSM700 or LSM900 with airyscan 2 were used. Images were then processed using Fiji (Schindelin *et al*, 2012)

### Antibodies

Primary antibodies: CASPR (1:500, Antibodies Inc., 75-001, RRID: AB_2083496), JUN (1:500, Cell Signalling Technology, 9165, RRID:AB_2130165), MBP (EMD Millipore, 1:1000, AB9348, RRID: AB_2140366), MPZ (1:500, Aves Labs, PZO, RRID: AB_2313561), NFL (1:500, Abcam, ab72997, RRID: AB_1267598), PRX (1:500, kind gift from Peter Brophy), SOX10 (1:100, R&D Systems, AF2864, RRID:AB_442208).

Secondary antibodies: Donkey anti-goat IgG (H+L) Alexa Fluor 488 (1:500, Thermo Fisher Scientific, A11057, RRID:AB_2534102), Goat anti-mouse IgG (H+L) Alexa Fluor 488 (1:500, Thermo Fisher Scientific, A11001, RRID:AB_2534069), Goat anti-mouse IgG (H+L) Alexa Fluor 647 (1:500, Thermo Fisher Scientific, A-21235, RRID:AB_2535804), Goat anti-rabbit IgG (H+L) Alexa Fluor 568 (1:500, Thermo Fisher Scientific, A11011, RRID:AB_143157), Donkey anti-chicken IgY (H+L) Alexa Fluor 488 (1:500, Jackson ImmunoResearch, 703-545-155, RRID:AB_2340375), Donkey anti-chicken IgY (H+L) Alexa Fluor 647 (1:500, Thermo Fisher Scientific, A78952, RRID:AB_2921074).

### Electron Microscopy

After removal of the microfluidic chamber, the orientation of the coverslips was marked, and cells were fixed overnight at 4°C in 2% glutaraldehyde/2% formaldehyde in 0.05 M sodium cacodylate buffer pH 7.4 containing 2 mM calcium chloride. After washing 5x with 0.05 M sodium cacodylate buffer pH 7.4, samples were osmicated (1% osmium tetroxide, 1.5% potassium ferricyanide, 0.05 M sodium cacodylate buffer pH 7.4) for 3 days at 4°C. Samples were washed 5x in deionised water (DIW) and treated with 0.1% thiocarbohydrazide/DIW for 20 minutes at RT in the dark. After 5 further washes in DIW, samples were osmicated a second time for 1 hour at RT (2% osmium tetroxide/DIW). After washing 5x in DIW, samples were block stained with uranyl acetate (2% uranyl acetate in 0.05 M maleate buffer, pH 5.5) for 3 days at 4°C. Samples were washed 5x in DIW and then dehydrated in a graded series of ethanol (50%/70%/95%/100%/100% dry) and 100% dry acetonitrile, 3x in each for at least 5 minutes. Samples were infiltrated with a 50/50 mixture of 100% dry acetonitrile/Quetol resin (12 g Quetol 651, 15.7 g NSA, 5.7 g MNA, all from TAAB) overnight, followed by 5 days in 100% Quetol resin with 0.5 g BDMA (TAAB), exchanging the resin each day. Aclar^®^ coverslips were placed on top of round polyethylene cups, with cells facing the resin. Samples were cured at 60°C for 3 days, and coverslips removed. The required section plane was marked on the block, and smaller sample blocks were cut from the resin using a hacksaw and mounted on resin stubs. Thin sections (∼ 70 nm) were prepared using an ultramicrotome (Leica Ultracut E) and collected on bare Cu TEM grids or Cu/carbon film grids. Samples were imaged in a Tecnai G2 TEM (FEI/Thermo Fisher Scientific) run at 200 keV using a 20 μm objective aperture to improve contrast. Images were acquired using an ORCA HR high resolution CCD camera (Advanced Microscopy Techniques Corp, Danvers USA).

### Live imaging of cocultures

Microfluidic chambers were placed on a Zeiss LSM 900 confocal microscope equipped with a temperature-controlled chamber at 37°C 5%CO_2_. Multiple areas of interest were selected for each microfluidic chamber and imaged every 10 minutes for up to 48 hours. To quantify number of SCs with fragments, each cell was defined as a region of interest and checked for the presence of mCherry positive fragments at all timepoints.

### Statistical Analysis

Results are shown as mean ± SEM. Statistical significance was estimated by Student’s t test or 2-way ANOVA. p < 0.05 was considered statistically significant. Statistical analysis was performed using GraphPad Prism software (version 9.5.0).

### Step by step dissociated mouse myelinating SC/DRG compartmentalised coculture protocol

#### Dissociated DRG neuron culture

1 Aclar^®^ plastic coverslip preparation (Electron microscopy sciences 50425-10)

- Cut into 40×22 mm pieces
- Autoclave in glass dish

2 Day 1
  2.1 Preparing the dishes

- Make a 750 μl drop of 0.5 mg ml^-1^ PLL (Merck - P1274) in a dish and place one Aclar^®^ coverslip on this drop
- Leave to coat overnight at room temperature (RT)

2.2 DRG dissection

- Dissect as many DRGs as possible from E14 mice:
- Place the embryo in a dish containing ice-cold L15 (Thermo Fisher - 11415049) in a sterile dissection hood
- Lay the embryo on its side and remove head and ventral part of the embryo (remove skin and internal organs)
- Remove any remaining tissue in front of the vertebral column
- Place vertebral column ventral side up and use micro-dissecting scissors or forceps to cut/crush through vertebral column
- Tease apart the right and left side of the vertebral column to expose spinal cord and DRGs
- Gently pull and remove the spinal cord and move it to a 35 mm dish containing Hibernate media (see 3.2) + NGF (Thermo Fisher - 13257019)
- Pluck off DRGs using forceps
- Store overnight at 4°C

3 Day 2
  3.1 Preparing the PLL-coated Aclar^®^ coverslips

- Wash coverslip with sterile ultrapure H_2_O (resistivity 18.2 MΩ·cm at 25°C)
- Airdry coverslips and move to a 60 mm dish
- Place microfluidic chamber on top of coverslip using sterilised forceps (kept in 100% EtOH)
- Check chamber attachment under microscope until there are no visible air bubbles between the chamber and the coverslip
- Place 3x 60 mm dishes in an upside down 200 mm dish

3.2 Matrigel^®^ Coating

- Thaw growth factor depleted Matrigel^®^ (Corning - 356231) on ice and dilute 1:200 in cold DMEM high glucose (4500 g dl^-1^ glucose; Thermo Fisher - 41966029)
- For both top and bottom compartments, pipette 150 μl into one well making sure it flows through the chamber into the other well: make sure to pipette forcefully in one fluid motion right at the channel entrance

- Take whole volume and pipette into other well to encourage flow
- Leave for at least 1 hour
- Add 35 mm dish full of water to maintain humidity and stop the chambers from drying out

3.3 Dissociation

- You will need approximately 10 DRGs per chamber
- Warm up 0.025% Trypsin (Merck - T9201) dissolved in Ca^2+^ and Mg^2+^ free PBS (Merck)
- Make up DRG media (section 4.3) and prepare DRG media + NGF (2.5ml per 10 ganglia/chamber, includes topping up next day)
- Transfer approximately 40 ganglia into 2.5ml of trypsin in a 15ml falcon
- Leave for 30 minutes at 37°C
- Warm up collagenase: 682 U ml^−1^ Collagenase (Worthington Biochemical Corporation - LS004176) in Ca^2+^ and Mg^2+^ free medium (10% Krebs Solution (133 mM NaCl, 177 mM KCL, 1.75 mM NaH2PO4, all Merck), 1% MEM Non-Essential Amino Acids Solution (Thermo Fisher), 0.5% Phenol Red Solution (Merck), 0.2% NaHCO3 (Thermo Fisher), 0.2% Glucose (Thermo Fisher))
- Transfer ganglia into 600 μl of collagenase in a 1.5 ml eppendorf (transfer only ganglia, not trypsin)
- Mix liquids but not cells
- Leave for 30 minutes at 37°C
- Prepare one 1 ml DRG medium in a 15 ml falcon per eppendorf of DRGs
- Transfer each eppendorf of DRGs to 1 ml DRG medium in a 15 ml falcon
- Triturate with P1000 pipette very gently - do not over dissociate, move any lumps of tissue left to a separate tube to triturate further
- Centrifuge for 5 minutes at 1200 rpm
- For lentiviral infection: resuspend in 1 ml DRG medium + NGF
- For plating: resuspend in DRG medium + NGF (1 μl per ganglion) and pool tubes

3.4 Lentiviral infection of DRGs

- Lentiviruses stored at −80°C and thawed on ice (can re-use lentivirus once but double concentration as viral copies reduce by 50% every freeze-thaw cycle)
- Add virus to DRGs resuspended in 1 ml of DRG medium + NGF
- Multiplicity of infection (MOI) of 2-10
- Leads to approximately 100% transduction
- Centrifuge for 1 hour at 400 rpm
- Resuspend pellet in 1 μl per ganglion DRG medium + NGF + virus (same amount)
- Proceed to plating

3.5 Plating

- Remove Matrigel^®^ from chamber and replace with 100 μl of DRG medium per compartment
- Remove all medium on plating side (leaving just medium in the channel to avoid air bubbles)
- load 10 μl cells into the top channel
- check underneath microscope and if cells look sparse, load another 5-10 μl of cells from the other side
- allow 4 hours for cells to attach
- top up with 100 μl added to top compartment: make sure to pipette from well to well to keep volume the same and reduce flow
- check that the media level is higher in the bottom compartment - if not, add one or two drops to establish volume difference

4 Day 3
  4.1 Topping up

- For square chambers (SND150, Xona Microfluidics^®^) top up wells (200 μl in top compartment and 300 μl in bottom compartment)
- For round chambers (RND150, Xona Microfluidics^®^) top up wells (75 μl in top compartment and 150 μl in bottom compartment)
- check that the media level is higher in the bottom compartment - if not, add one or two drops to establish volume difference
- Add anti-mitotic agent cytosine arabinoside (Ara-C, Merck - C6645) to top compartment: final concentration 10^−5^ M, add half of the volume per well

5 Changing Media

- Change medium on Mondays, Wednesdays, and Fridays
- Make sure to pipette one drop per well into alternating wells to reduce flow
- Make sure to top up the water in the dish to stop chambers drying out

6 Day 10

- After maintaining DRGs in Ara-C for 7 days change media to DRG + NGF without Ara-C
- Non-neuronal cells will not return after this stage
- Wait for axons to extend to the wells before seeding SCs AND for Ara-C to be removed 3 days prior

### Schwann Cell Culture

- Dissect P3-5 sciatic nerves and brachial plexuses from mice and start SC culture within 3-5 days of DRG dissociation
- After 3 days of Ara-C purification, trypsinise cells and infect with lentiviruses in suspension
- Expand SCs on 60mm PLL/laminin coated dishes until needed for coculture

1 Preparing dishes
  1.1 PLL Coating

- Prepare a 0.2 mg ml^-1^ solution of PLL and coat 35 or 60 mm sterile dishes overnight
- Remove PLL (can be re-frozen and used 3 times)
- Wash 3 times with sterile ultrapure H_2_O (resistivity 18.2 MΩ·cm at 25 °C)
- leave dishes to air dry, store at RT

1.1 Laminin Coating

- Dilute the stock solution of laminin (Merck - L2020) in low glucose DMEM (1000g/dl; Thermo Fisher - 21885025) to a final concentration of 10 μg ml^-1^ (1/100 dilution)
- Add the solution to the dish
- Leave for at least 1 hour at 37°C
- Remove laminin immediately prior to plating cells (can be reused 3 times) and do not let dishes dry (add media)

1 Mouse SC Purification

- Make 20 ml of DMEM low glucose (with 1/100 Pen/Strep (Thermo Fisher - 15140122)) + 5% HS (Thermo Fisher - 16050130) and warm to 37°C
- Prepare 2x 60 mm tissue culture dishes of L15 and place on ice
- Dissect out sciatic nerves and brachial plexuses and place in ice-cold L15
- De-sheath the nerves and transfer to a separate dish containing L15
- Place 100 μl trypsin and 100 μl collagenase (per 2 animals) in a 35 mm dish: 2 mg ml^-1^ Trypsin (Merck - 85450C) and 682 U ml^−1^ Collagenase in Ca^2+^ and Mg^2+^ free medium
- Transfer the nerves into the trypsin/collagenase and incubate at 37°C for 45 minutes
- Triturate nerves with a P1000 and then with a P200
- Stop the digestion by adding an excess 2 ml low glucose DMEM + 5% HS
- Transfer the cell suspension to a 15 ml centrifuge tube
- Centrifuge at 1000 rpm for 10 minutes
- Resuspend the cell pellet in 2 or 4 ml of low glucose DMEM + 5% horse serum and transfer to the 35 or 60 mm laminin-coated dish
- Add Ara-C to a final concentration of 10^−5^ M and culture for 3 days to eliminate fibroblasts
- After 3 days, replate SCs

1 Mouse SC Expansion

- Mouse SCs proliferate on PLL/laminin coated dishes in the presence of
- low serum (0.5% HS)
- βNRG1 (R&D Systems - 396-HB-050)
- a low cAMP signal (50-100 μM dibutryl-cAMP (dbcAMP; Merck - D0627) or 2

#### μM forskolin (Merck - 344270))

- Laminin coat 60 mm PLL coated plates
- Wash cells 2x with PBS
- Trypsinise Ara-C purified SCs using 1 ml of 6% 2 mg ml^-1^ Trypsin (Merck - 85450C) in Versene (0.02% EDTA (Thermo Fisher - D/0700/53) + 0.5% Phenol Red in PBS) for up to 5 minutes
- Stop the reaction by adding 2 ml (or more) of low glucose DMEM + 5% HS
- Transfer the cell suspension to a 15 ml centrifuge tube
- Centrifuge at 1000 rpm for 10 minutes
- Pre-plate the cells to eliminate fibroblasts if needed:
- resuspend pellet in 10 ml of low glucose DMEM + 5% HS or defined medium (DM, table 1) with 0.5% HS (if they have already been expanded in DM)
- add to an uncoated 90 mm tissue culture dish (no PLL, no Laminin) for 2-3 hours
- fibroblasts will sit down and attach
- wash the dish well
- collect medium
- centrifuge at 1000 rpm for 10 mins
- Resuspend in SC expansion medium (DM + 0.5% HS + 10 ng ml^-1^ βNRG-1 + 2 μM forskolin) and plate or lentivirally infect prior to expanding
- Change medium every 3 days
- Split cells when 80% confluent
- Do not passage cells more than 3 times as mouse SC tend to quiesce after this
- Lentiviral Infection of Mouse SCs
  - Pre-plate cells as above and then lentivirally infect prior to expanding
  - Resuspend in 1 ml of DM + 0.5% HS
  - Calculate the number of cells using a haemocytometer
  - Lentiviruses stored at −80°C and thawed on ice
  - Add virus (MOI of 200-500)
  - Centrifuge for 1 hour at 400 rpm
  - Resuspend pellet the pellet without changing the medium as you want to keep the virus
  - Plate SCs
  - Change medium after 24 hours to fresh SC expansion medium
  - Proceed with normal expansion
  - Quiescing SCs
    - The day before they are used in an experiment, SC cell cycles are synchronised by quiescing them
    - Change media to DM + 0.5% HS WITHOUT any forskolin or neuregulin

### Coculture

- Seeding SCs

- Medium change DRG cells side into DRG/SC medium + NGF (section 4.4)
- there are roughly 400,000-600,000 SCs in a confluent 60 mm dish (200,000 if 50% confluent)
- Trypsinise SCs with 1 ml of 6% 2 mg ml^-1^ Trypsin in Versene for max 5 minutes at 37°C
- Stop the reaction with DMEM low glucose + 5% HS
- Centrifuge for 10 minutes at 1200 rpm
- Resuspend in DRG/SC medium + NGF to achieve a concentration of 30,000 cells per 10 μl media (3,000,000 ml^-1^)
- Remove almost all the medium from bottom compartment
- Load 30,000 cells by pipetting at the entrance of the channel
- Check underneath microscope and if cells look sparse, load another 5-10 μl of cells from the other side
- After 4 hours, top up medium to normal levels
- Change medium on Mondays, Wednesdays, and Fridays
- Inducing Myelination

- Allow 7 days for SCs to align and proliferate before inducing myelination
- Keep top compartment in DRG/SC medium + NGF
- Change bottom compartment to axon only media, keep in DRG/SC + NGF (for aligned SC), or myelination media
- Change medium on Mondays, Wednesdays, and Fridays

### Media

1. Supplement stocks

- NGF: 100 μg ml^-1^, Thermo Fisher Mouse NGF 2.5S Native Protein (13257019)
- Forskolin: 10 mM in ethanol, Merck, Coleus forskohlii - CAS 66575-29-9 – Calbiochem
- Neuregulin: 10 μg ml^-1^ in in PBS 1% BSA, Recombinant Human NRG1-beta 1/HRG1-beta 1 EGF Domain Protein, RD Systems, 396-HB-050
2. Hibernate DRG medium

- for 50 ml of media:

48 ml Hibernate E (Thermo Fisher - A1247601)
1 ml B27 (2%; Thermo Fisher - 17504044)
500 μl Pen/Strep
500 μl L-Glutamine (Thermo Fisher - 25030081)
- on day of use add NGF to a final concentration of 33 ng ml^-1^ (1/1000)
3. DRG medium

- for 50 ml of media:

48.5 ml DMEM (high glucose)
1 ml B27
500 μl Pen/Strep
- on day of use add NGF 1/3000
4. DRG/SC medium

- for 50 ml of media:

24.5 ml DMEM (high glucose)
24.5 ml DM
1 ml B27
500 μl Pen/Strep
- on day of use add NGF 1/3000
5. Defined medium (DM; Jessen et al., 1994)

- prepare medium according to table 1 and store at 4°C
6. Axon only medium

- for 50 ml of media:

24.5 ml DMEM (high glucose)
24.5 ml DM
1 ml B27
500 μl Pen/Strep
- on day of use add

NGF 1/3000
Forskolin 1/1000
NRG 1/1000
7. Myelination medium

- for 50 ml of media:

24.5 ml DMEM (high glucose)
24.5 ml DM
1 ml B27
500 μl Pen/Strep
- on day of use add

Matrigel^®^1/100 (you can make a 5 or 10 ml stock with Matrigel^®^ added)
NGF 1/3000
Forskolin 1/1000
NRG 1/1000

**Table 1:** Defined medium

|  | Reagent | Source | Final Concentration |
| --- | --- | --- | --- |
| 50 ml | Hams F12 | Thermo Fisher - 31765035 | 47% |
| 50 ml | DMEM (low glucose) | Thermo Fisher - 21885025 | 47% |
| 2 ml | L-Glutamine | Merck - 25030081 | 2 mM |
| 1 ml | Penicillin Streptomycin | Thermo Fisher | 100 μM |
| 1 ml | 3.5% Bovine Serum Albumin | Merck - A7979 | 0.035% |
| 1 ml | 1.6 mg ml<sup>-1</sup> Putrescine | Merck - P5780 | 16 μg ml<sup>-1</sup> |
| 1 ml | 10 mg ml<sup>-1</sup> Transferrin | Merck - T1283 | 100 μg ml<sup>-1</sup> |
| 100 μl | 400 μg ml<sup>-1</sup> L-Thyroxine (T4) | Merck – T1775 | 400 ng ml<sup>-1</sup> |
| 100 μl | 0.06 mg ml<sup>-1</sup> Progesterone | Merck - P0130 | 60 ng ml<sup>-1</sup> |
| 100 μl | 5.7 mg ml<sup>-1</sup> Insulin | Merck - I9278 | 10<sup>-6</sup> M |
| 77 μl | 0.05 mg ml<sup>-1</sup> Dexamethasone | Merck - D4902 | 38 ng ml<sup>-1</sup> |
| 10 μl | 0.101 mg ml<sup>-1</sup> L-Thyronine (T3) | Merck - T6397 | 10.1 ng ml<sup>-1</sup> |
| 10 μl | 1.6 mg ml<sup>-1</sup> Selenium | Merck - S5261 | 160 ng ml<sup>-1</sup> |

- To wells, also add 50 μg ml<sup>-1</sup> L-Ascorbic Acid (1/100 from stock; Merck - A4544), make up stock (in H<sub>2</sub>O) fresh each time and protect from light

## Acknowledgements

We thank Karin Müller, Martin Lenz, and Filomena Gallo at the Cambridge Centre for Advanced Imaging for assistance with microscopy. We thank Aviva Tolkovsky and Alex Clark for technical suggestions. We thank Peter Brophy for the gift of the periaxin antibody. We thank Claire Jacob for helpful suggestions. For the purpose of Open Access, the author has applied a CC BY public copyright licence to any Author Accepted Manuscript version arising from this submission.

---

**Supplemental video 1: Axonal fragments are present in Schwann cells after axotomy.** 48-hour confocal live imaging of mCherry-labelled axons (magenta) and GFP-labelled SCs (green). Initially, intact axons are present, that then swell and break into large pieces, surrounded by SC processes resembling the constricting actin spheres shown by Vaquié et al., 2019. Larger axonal fragments are broken into smaller pieces and are transported along SC projections towards the perinuclear cytoplasm. Cultures were imaged every 10 minutes for 48 hours. 12 frames per second.

